# Optimal spatial monitoring of populations described by reaction-diffusion models

**DOI:** 10.1101/2021.05.21.445122

**Authors:** Nicolas Parisey, Melen Leclerc, Katarzyna Adamczyk-Chauvat

## Abstract

Using spatialised population measurements and related geographic habitat data, it is quite feasible nowadays to derive parsimonious spatially explicit population models and to carry on their parameter estimation. To achieve such goal, reaction-diffusion models are fairly common in conservation biology and agricultural plant health where they are used, for example, for landscape planning or epidemiological surveillance. Unfortunately, if the mathematical methods and computational power are readily available, biological measurements are not. Despite the high throughput of some habitat related remote sensors, the experimental cost of biological measurements are, in our view, one of the worst bottleneck against a widespread usage of reaction-diffusion models. Hence, in this paper, we will recall some classical methods for optimal experimental design that we deem useful to spatial ecologist. Using two case studies, one in landscape ecology and one in conservation biology, we will show how to construct *a priori* experimental design minimizing variance of parameter estimates, enabling optimal experimental setup with pre and post hoc filtering for accommodating additional constraints.

## 1 Introduction

Both empirical and theoretical studies have well established that population spread is an essential ecological process for understanding most of the observed population dynamics. Consequently, since the last two decades spatial ecology has become central either in theoretical ecology or in empirical approaches. Hence, population management for their control, capture or protection, has started to be thought at large spatial scales that match the inherent dispersal capacity of the considered species. Populations can be identified and monitored using various strategies (e.g. tracking, trapping). However, despite the development of new technologies and devices for ecological monitoring (e.g. imaging sensors for animal detection [1], autonomous Unmanned Aerial Vehicle [2]), the survey of many species on large areas remains challenging, costly and empirically guided.

The growing interest for space in ecology has been accompanied by a proliferation of models and statistical methods for analyzing and predicting the spatial distribution of populations. Nowadays, existing statistical methods allows one to infer the dynamics of spreading populations from noisy and sparse data. However, statistical inference of spatio-temporal models from common monitoring data can be subject to practical identifiability issues and is often associated with an important uncertainty on parameters estimates. Thus, given the inherent difficulty of collecting spatial data on most population, model-based forecasts are still associated with an important uncertainty, which could theoretically be reduced by more dense and efficient monitoring strategies.

When planning a survey of population over a large area, one has to make some choices giving financial and human constraints: which observational method (transects, traps), where and when to start the survey, the number and times for repeating the survey. These questions can either be addressed empirically or using model-based techniques that can be organized into three major groups of methods [3]. The first group transform the problem into a state-estimation one by augmenting the state vector [4]. The second group is based on random fields theory [5] whereas the third group corresponds to the classical theory of the design of experiments [3]. We can also point out a fourth group that consists in adopting an heuristic to find the best sampling strategies among tested situations [6]. Based on either of these groups, methods for designing ecological sampling has already been addressed by several authors [7, 8] but it has not yet percolated in the modellers community and is still seldom considered in population ecology. The use of optimal design of experiments is perhaps one the lesser known method [9] whereas it is well developed for optimal sensor placements in spatially-distributed systems with non-linear dynamics models [3]. It corresponds to an entire branch of statistics that provides criterion measuring the amount of information about unknown model parameters carried out by the observed data. Here, “experiments” is taken *sensus lacto* and refers to controlled observations of populations within their living areas that are planned under statistically optimal conditions given the constraints of available resources.

In this study we use the optimal design of experiments to tackle the question of ecological monitoring of populations described by reaction-diffusion models. This type of partial differential equations originally emerged in chemistry for analyzing the change in space and time of chemical substances before becoming one of the most important mathematical model for the study of spreading dynamical processes in biology and ecology. Here, we consider reaction-diffusion equations that provides a parsimonious description of a spreading population in a domain Ω included in ℝ ^2^ :

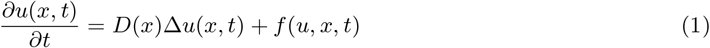

where *u*(*x, t*) is the population density at time *t* and location *x* ∈ Ω, *D*(*x*) corresponds to the dispersion rate at *x*, Δ*u*(*x, t*) is the Laplace operator that describes the random movement of individuals, and *f* (*u, x, t*) is a general reaction function accounting for local birth, death and interactions within the spreading population. To fully describe the system, one needs to define some boundary conditions on Ω as well as initial conditions *u*_0_(*x*) = *u*(0, *x*).

In addition to their ability to capture the essential observed patterns of spreading populations, reaction-diffusion systems have been well studied by mathematicians for a long time. Thus, though the mathematical and the numerical analysis of some reaction-diffusion equations still challenge mathematicians, several systems are now well characterized and understood. The most popular one is the Fisher-KPP system, that applies in ecology, for which *D*(*x*) = *D* and the reaction term accounts for a limiting carrying capacity of the environment *f* (*u, x, t*) = *u*(1 − *u*). Parameter estimation of mechanistic PDE models from noisy observational data, obtained during common population monitoring strategies, can be performed using different statistical methods. In any case, one needs to link the mechanistic PDE model that describes the changes in space and time of the population density with a probabilistic model describing the observation process. This approach refers to the physical-statistical or mechanistic-statistical models that are specific instances of the more general hierarchical state-space model framework [10, 11, 12]. Such approach has been successfully used for inferring spatio-temporal processes on large areas for invasive and beneficial insects [13, 14, 15], plant diseases [16] or aquatic and terrestrial mammal species [17]. As illustrated by [8], the optimal design of experiments can be applied on mechanistic-statistical systems for ecological monitoring, but is yet poorly considered by both modellers and ecologists collecting spatio-temporal data.

In this article we first introduce the main concepts of the optimal design of experiments applied on mechanistic-statistical systems for population dynamics. Then, we consider two example populations whose spatio-temporal dynamics is described by reaction-diffusion equations, but monitored with different strategies : beneficial insects for agriculture that are monitored using Barber pitfall traps at various locations within the landscape [13], and invasive wild horses that are counted during aircraft transects [18]. Assuming that the more realistic degree of freedom in the existing monitoring designs is the choice of the locations of observations we consider static designs, where locations of sampled populations are fixed before the start of the survey and don’t change in time, and focus on the spatial aspect of monitoring. We use D-optimal designs that define optimal locations for population observation and show how this framework can help to design ecological monitoring strategies over large areas under identical constraints. We finish the paper by discussing the use of the optimal design of experiments framework to improve the monitoring of populations over large areas, especially to reduce the cost of sampling and support the link between modelling and empirical studies.

## 2 Mechanistic-statistical model for population dynamics

We consider a mechanistic-statistical approach that combines a reaction-diffusion system which describes essential spatio-temporal dynamics of a population with an observational equation describing the stochastic process leading to detection and enumeration of individuals at location *x* ∈ Ω and time *t* (as in [11]). However, following the framework used in control system analysis or optimal design for parameter identification [3], we decompose the observation process into two parts : 1) the measurement process which takes into account factors that affect the measurement of population density (e.g. equipment features, operator behaviour, environmental effect), and 2) the data process that links population data with both the ecological and measurement processes. Then, given some boundary conditions on Ω the spatially distributed system is described by the following hierarchical system :

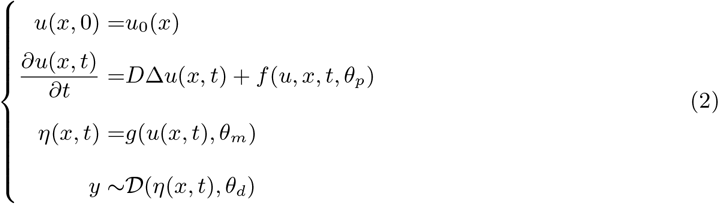

where *η*(*x, t*) is the measurement process determined from population density *u*(*x, t*) with function *g*(.), and the data process corresponds to the draw of population data *y* from a general probability distribution *𝒟*. Depending on the considered system, parameters that needs to be estimated, or identified, from the monitoring data *y* can occur in the population process (*D, θ*_*p*_), the measurement process (*θ*_*m*_) and the data process (*θ*_*d*_). The set of unknown parameters is thus given by *θ* = (*D, θ*_*p*_, *θ*_*m*_, *θ*_*d*_). Albeit initial conditions *u*_0_(*x*) of reaction-diffusion systems may also be estimated from noisy data [16, 11], in this study we assume that *u*_0_(*x*) is known and fixed. Parameter estimation can be achieved using maximum likelihood estimation (see next section). It generally involves a numerical optimization that requires the simulation of the reaction-diffusion system with a suitable numerical method [19, 20].

## 3 Optimal design

For monitoring a population, the questions that we should address at first are where to look at the population and when to observe the targeted population ? Though the time scheduling of the monitoring is often constrained by numerous factors, the choice of spatial locations generally offers more degree of freedom. In the following, we assume that the dates of the monitoring design (or experiment *sensus lato*) is scheduled in advance and also that the sensors locations will not change in time. This simplifies the problem of defining optimal monitoring designs and thus an optimal spatial distribution of observational locations over the survey area.

This question can be addressed through experimental design theory, a branch of applied statistics introduced by Sir Ronald Fisher in his seminal book edited in 1935 [21]. The purpose of this section is to introduce some key-notions of experimental design theory, essential for understanding its application on the use cases presented in section 4. For a complete description of the statistical framework the reader can refer to Silvey [22], Atkinson [23] or Walter and Pronzatto [24]. We start by introducing the key notions of optimal design and then we gave their interpretation for the mechanistic-statistical model presented in equation 2. We indifferently use the terminology inherited from the design of experiments theory. Therefore, the experiments will refer to the survey of the populations and sensors corresponds to any monitoring device or a visual inspection.

Let us consider a random variable *y* with a probability distribution depending on: (1) a vector of real variables *x* that can be chosen by the experimenter, (2) a vector of parameters *θ*, supposed to be fixed and unknown for the experimenter. We assume that *x* belongs to the set *𝒳 ⊂* ℝ^*r*^ and that *θ* belongs to a parametric space Θ *⊂* ℝ^*p*^. The variables composing *x* are called control variables. Let us suppose that for a given *x* and *θ* the distribution of *y* is given by a probability density function *p*(*y*|*x, θ*). The experimenter is allowed to take *N* independent observations on *y* at vectors *x*_1_, …, *x*_*N*_ chosen from the set *𝒳*. The set **x** = *{x*_1_ … *x*_*N*_ *}* will be referred as *N* -observation design. The question, primarily, is how to select the design **x** ?

The criterion of choice depends on the purpose of the experiment. Here our primary interest is in estimating the parameter *θ* from the experimental data. Let us denote by *<u>x</u>* a vector (*x*_1_, …, *x*_*N*_) and let *<u>y</u>* be a vector of values of *y* taken at *x*_*i*_. The log-likelihood function of *θ* is defined as:

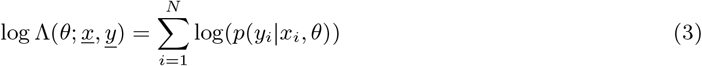

The maximum likelihood estimate of *θ* maximizes log Λ over Θ:

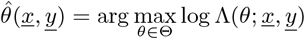

Under some regularity conditions on the family of densities *{p*_*θ*_ : *θ* ∈ Θ*}* the estimate 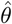 is asymptotically normal as the sample size *N* tends to infinity (see [25] for instance). Moreover, the asymptotic variance of 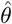 reaches its lower bound given by Cramer-Rao inequality and is equal to the inverse of the Fisher Information Matrix, defined as:

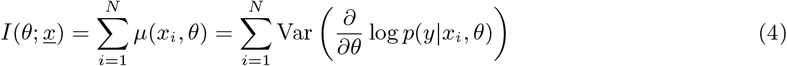

where *µ*(*x, θ*) is the information matrix for an observation on *y* taken at *x*. Roughly speaking an optimal design **x**_*\**_ makes the variance of 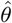 “as small as possible” or, alternatively, makes the Fisher matrix “as large as possible”. To be more precise we seek for an **x**_*\**_ that maximizes some real-valued function *ϕ* of *I*(*θ*; *<u>x</u>*). We assume here that *ϕ* is homogeneous and concave.

We consider the design with *n* distinct vectors *x*_1_, …, *x*_*n*_ replicated *r*_1_, …, *r*_*n*_ times, where 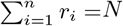. We assign to **x** a discrete probability distribution *ξ*^*n*^ which puts the probabilities *ω*_*i*_ = *r*_*i*_*/N* at *x*_*i*_. Consequently, the Information Matrix (4) can be rewritten as :

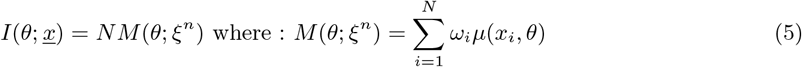

As a criterion function *ϕ* is homogeneous then optimizing *ϕ*(*I*(*θ*; *<u>x</u>*)) amounts to optimizing *ϕ*(*M* (*θ*; *ξ*_*n*_)) over *ξ*^*n*^. An optimal design 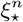 is thus defined as:

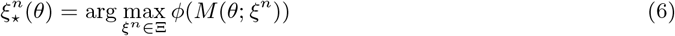

where the matrix *M* is assumed to be non-singular.

### D-optimal design

The choice of the optimality criterion relies on the final purpose of statistical analysis. The experimenter may wish to improve the precision of 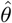 or to reduce the variance of the predicted values of *y* or again to better discriminate between the candidate models. The main optimality criteria and the corresponding functions *ϕ* can be found for example in [24]. Because of its versatility of purpose, one of the mostly used criterion is the D criterion leading to a D-optimal design, maximizing the logarithm of the determinant of the matrix *M* :

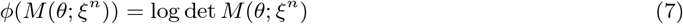

A D-optimal design aims at minimizing the volume of confidence ellipsoid for model’s parameters. Consequently the statistical treatment of experimental results become more efficient. For example, it help to assess if a model’s parameter, and its linked hypothesis, has a non negligible effect on a population dynamics e.g. is a parameter different from zero. The D-optimal design also helps strengthen parameters comparison. For example it’s usually important to estimate which habitat is the best (or worst) for a given species e.g. to significatively rank their corresponding growth rates. Finally, this design can also lead to minimizing the maximum variance of the predicted values [26].

### Exact and approximate design

When the set of the matrices defining a domain of (7) is discrete, the design is said to be exact.The problem of finding an exact optimal design 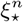 could be numerically challenging, especially for the large values of *N*. The solution proposed in the theory of optimal design is to calculate the optimum of *ϕ* over the extended domain and to look for the discrete N-observational design which is “close” to the optimum. The definition of the matrix *M* can be extended by considering the set of all probability distributions on *𝒳* in place of the discrete distributions *ξ*^*n*^:

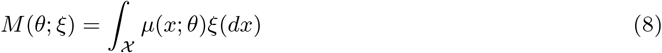

where *ξ* is a probability distribution on *𝒳*. Then the approximate solution of the initial optimization problem can be calculated. An optimal (continous) approximate design *ξ*_*\**_ is defined as:

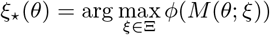

for the matrix *M* given by the formula (8).

### Average and local optimal design

An optimal design for non-linear model depends on the unknown parameter value and is referred to as a local design. Different methods are proposed to “remove” the dependence from *θ*. One possible approach is to assume that *θ* is a random variable following a known distribution *π* and to maximize the expected value of the criterion, calculated with respect to *π*. Consequently, an on-average exact design maximizes:

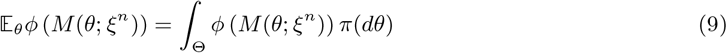

### Fisher Information Matrix for generalized regression model

According to the definition given in [27], generalized regression model is specified by the probability density *p* that depends on *x* and *θ* through the *k* dimensional regression function *η*(*x, θ*):

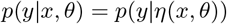

Consequently, the Fisher Information Matrix *µ*(*x, θ*) for a single observation on *y* can be rewritten as :

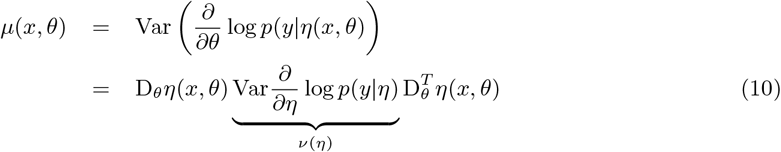

where D_*θ*_*η*(*x, θ*) is a *p* × *k* matrix of the first derivatives of the regression function with respect to *θ* and *ν*(*η*) is a *k* × *k* Fisher Information Matrix for the reparametrized model. The matrix *ν*(*η*) is called the elemental information matrix and is known for many useful probability densities.

### Design settings for spatial population monitoring

We assume that the targeted population is observed at *N* locations *x*_*i*_, …, *x*_*N*_. For each location, the measurement is repeated *T* -times. The time measurements are fixed in advance and common for all the locations whereas the set of the locations is solution to the optimal design problem. The measurement taken at a location *x*_*i*_ at a time moment *t* is denoted by *y*(*x*_*i*_, *t*). Typically *y*(*x*_*i*_, *t*) is either a population count or its suitable transform. In terms of mechanico-statistical model introduced in section 2, *y*(*x*_*i*_, *t*) is the result of a data process, a random variable following a probability distribution *𝒟*. The distribution *𝒟* depends on a measurement process *η*(*x, θ*) governed by a partial differential equation and on a vector of parameters *θ*, related to all the levels of hierachical model (see eq. 2). The likelihood of *θ*, given the vector of observations 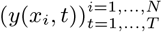 is given by equation 3. The optimal design problem adressed in this paper is to find an exact D-optimal on-average design 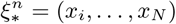 (implicitly *ω*_1_ = … = *ω*_*N*_ = 1). According to the short introduction given in this section, 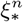 satisfies :

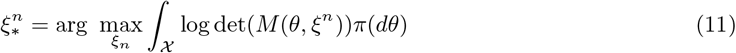

where *M* (*θ, ξ*^*n*^) is calculated according to equation 5 and 10. From this point on, we assume that the parameter *θ* follows a uniform distribution *𝒰* (*θ*_*min*_, *θ*_*max*_). In the following chapters, we illustrate the solution of optimal design 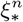 for two spatial monitoring examples.

## 4 Spatial population monitoring case studies

### 4.1 Agroecological case study

As a first example we consider the dynamics of *Poecilus cupreus*, a carabid beetle known for its weed seed consumption, within agricultural landscapes. This beneficial insect often complete its life cycle in a couple of months and is commonly monitored using pitfall traps that are picked up and replaced weekly. This species exhibits a single peak during its activity season and is known to react differently to semi-natural habitats, grasslands and cereal fields [28]. This leads to a reaction-diffusion model introduced in [13] with a birth rate decays parameter (*β*), to mimick the single pick, and a spatially heterogeneous growth rate *r*(*x*), to express the habitat dependencies. In [13], the population dynamics of carabids has been studied within a landscape of a few kilometer squared size. The dynamics of carabids was described by the following mechanistic-statistical model:

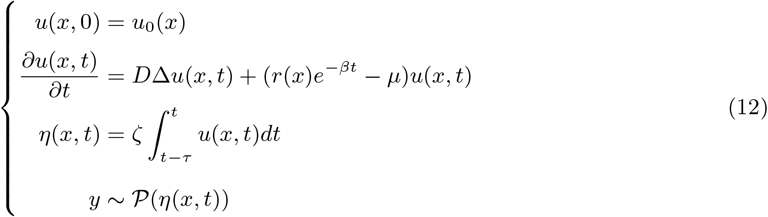

where *u*(*x, t*) is in this case the density of carabids, *β* describes the exponential speed at which the birth rate decays during the activity season, *µ* is the death rate (i.e.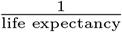), *r*(*x*) is an habitat specific growth rate such that :

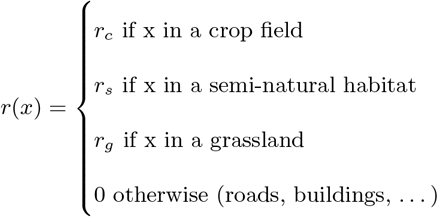

*ζ* is a scaling factor that allows the integration of the carabids density over the surface of a non-attractive pitfall trap from which count data is described by a Poissonian data process *𝒫* (.) with intensity *η*(*x, t*). The population process is parametrized by (*D, θ*_*p*_) with *θ*_*p*_ = (*β, µ, r*_*c*_, *r*_*s*_, *r*_*g*_) and the measurement process depends on *θ*_*m*_ = (*ζ*), that we considered known for a given type of traps. Hence the vector of unknown parameters were *θ* = (*D, β, µ, r*_*c*_, *r*_*h*_, *r*_*g*_). The parameters of the model were already estimated in [13] for an arbitrarily chosen set of pitfall traps. We used these results in order to fix the bounds of parameters *θ* in an on-average design. *θ*_*min*_ and *θ*_*max*_ were set so they were biologicaly relevant and coherent with previous estimations. All growth rates were explored so they give half to twice as many descendants, during a season of several months, as previously estimated. The diffusion coefficient varies between a population of individuals half to twice as fast. As *µ* is linked with life expectancy, it varies between a life twice as long and half as long. Finally, *β*, measuring birth decays during the activity season, varies between 0.9 and 1.1 times its nominal value. Details for all parameters values can be found in appendix A.1.

In this example, as described in [13], the experiment consisted in placing *N* = 24 pitfall traps in an agricultural landscape, owned by several farms, and sample them over the course of a season, in that case *T* = 9 times over the course of three months in 2010. The sampling design of this study was decided beforehand so that, first, an agricultural field was chosen then several traps (here three) were planted, spatially grouped, in the field. From now on, we will refer to this as a clustered design.

### 4.2 Conservation biology case study

As a second example we considered the dynamics of *Equus caballus*, an invasive feral horse that has detrimental effects to the ecosystem of the Australian Alps and whose management through aerial culling or trapping and mustering is under debate. This question has been addressed by [18] who used a two years aerial survey of feral horse populations, consisting in line transect data within the Australia Alps region [29]. They derived a reaction-diffusion model assuming horse populations are limited by density dependence in births via a logistic model, and that their movement through the landscape depends on the local horse density and, spatially heterogeneous, carrying capacity. We supplemented their spatio-temporal dynamics with a measurement process and a data process to obtain a full mechanistic-statistical model. The measurement process model the use of aerial transects to assess horse populations while the data process assumes normally distributed residuals with zero mean and variance *σ*^2^. This lead to a mechanistic-statistical system given by :

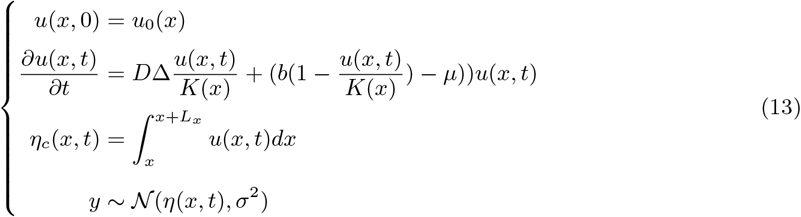

where *u*(*x, t*) is in this case the density of horses, *K*(*x*) the habitat-dependent carrying capacity, *b* the birth rate and *µ* the mortality rate. Population densities are monitored on transects of size *l* × *L*_*x*_ and drawn from a Gaussian distribution *𝒩* (.). We assume all transects have the same width *l* (from aerial line of sight) but that the length *L*_*x*_ will depend on each potential transect. The habitat-dependent carrying capacity was defined as :

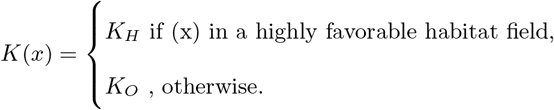

The population process is parametrized by (*D, θ*_*p*_) with *θ*_*p*_ = (*b, µ, K*_*H*_, *K*_*O*_) and the data process depends on *θ*_*d*_ = (*σ*^2^). In this second case study the question we addressed was to find the best *n* transects, over *N*. We chose to simulate a survey of the population performed 7 years after the original monitoring (i.e. 2021). In [18], the mortality rate *µ* was fixed and the carrying capacity *K*(*x*) was estimated independently of the population process. Hence, the vector of unknown parameters, for the conservation biology case, was *θ* = (*D, b*). One can note that *σ* is classically not part of such optimal design as it can be estimated by residual standard deviation. There was no previous estimations for *θ* but there were lower and upper boundaries, expressed as ‘low growth and dispersal’ and ‘high growth and dispersal’ scenarios in the original paper. We used them to set *θ*_*min*_ and *θ*_*max*_ used in the on-average exact design whereas for local design, we considered *θ*_*min*_. Details for all parameters values can be found in appendix A.1.

In this example, as described in [18], the experiment consisted in surveying *N* = 32 aerial transects, of different lengthes, one time, in the year 2014, to estimate wild horses populations in the autralian alps [29].

## 5 Criteria for comparing designs and numerical implementation

### 5.1 Performance assessment of designs

In optimal design, one can quantify the suboptimality of any given designs compared to the D-optimal design, as described by eq. 5 and 7, using the notion of the D-efficiency [30] which is defined as follows :

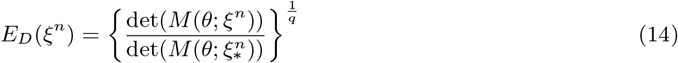

with *q* the number of parameters in *θ*. We can use this measure to compare our designs among themselves, or with any other designs, e.g. taken from the literature or even randomly drawn. It can also be of interest to visualize the design properties. Especially, one can review design positions on maps. One might also want to focus on the relationship between pair of parameters, using confidence ellipses [31] derived from the inverse of the Fisher matrix.

### 5.2 Evaluating designs according to a practical constraint

When planning an ecological monitoring the statistical properties of the design is only part of the decision making. One important factor for planning the monitoring in the two study cases considered here is the length of travelling that induces important costs. In order to integrate this component we consider the tour length of travelling salesman solutions [32] where one start at a relevant ‘base camp’, then tour a design before going back to camp. Formally, as defined in [33], we label the traps from 1, …, *n* and define :

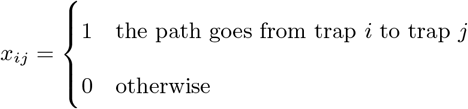

Taking *c*_*ij*_ > 0 to be the Euclidean distance from trap *i* to trap *j*, the travelling salesman problem (TSP) can be written as the following integer linear programming problem:

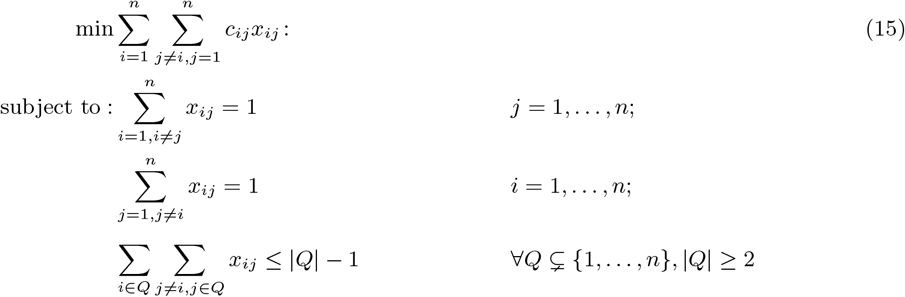

As one can see, the TSP minimize the tour length while ensuring each trap is arrived at from exactly one other trap, departed to exactly one other trap and that the solution returned is a single tour and not the union of smaller tours. For the second use case, involving lines instead of points, a heuristic is used instead of the problem 15. The tour length is roughly approximated by a weighted mean between the diagonal of the bounding box of the transects and their cumulated lengthes. Finally, the best compromises between statistically efficient designs and travelling lengths can be visualized on a Pareto front in the D-efficiency - tour length space [34]. Of course, we can note that performance assessment of an experimental design is not limited to the above examples.

### 5.3 Numerical implementation

We implemented different strategies to solve numerically the computationally demanding problems we worked on. As the computation of the on-average design (eq. 11) with a uniform distribution would be cumbersome, we relied on a sampling from a Latin Hypercube, a space-filing design [35], to ‘cover’ at best a domain with a limited number of sampled points.

Moreover, we noted that one useful feature that come at no cost with our designs is the ability to filter the set of *N* possible points (or transects) before searching for a solution. For example, in the agroecological case, we used a regular grid to select *M* possible trap locations within fields (*M* ≫*N*) while excluding road from pitfall trap’s possible positions.

In addition, in order to solve the information matrix that requires the derivatives of the deterministic process models we chose to numerically extract them ahead of solving the design, i.e. to do some memoization [36], which proved relevant for the task at hand. For performance, one could also rely on analytic, symbolic or automatic differentiation (adjoint code) if applicable.

Finally, we relied on an excursion algorithm [37], with multiple starts, for solving the designs. These algorithms take the particular form of the problem into account. Starting from a non-degenerated design (i.e. with positive determinant), at each iteration *k*, exchange one support point by a better one, in the sense of the D-optimality. Considering all *N \** (*M* − *N*) possible exchanges successively, retaining the best one among these each iteration, it converges to a local solution. Hence, we used multiple starts to get the best local solution as an approximation of a global solution.

Numerical calculations were performed in R [38]. Using a Runge-Kutta solver combined with the method of lines to solve the reaction-diffusion models [39] and some naive parallelization, it taked about an hour to solve an on-average design on a Intel^®^ Xeon^®^ E5 with 8 cores 8 Go of RAM. All code is available upon request.

## 6 Results

We reduced the size of the original landscapes, and thus the number of sampling points, to reduce the computational cost. For the agroecological case study, we used a virtual agroecosystem, of one km^2^, inspired by [11] with just six pitfal traps sampled over a season. For the conservation biology case study, we used a simplified habitat map, of several thousand km^2^, of a subpart of the australian alps (see supplementary materials of [18]), sampled one time, 7 years after the first experiment (hence in 2021). For this example, we reduced the number of surveyed transects to eight, chosen among the original thirty two, through D-optimality.

For each use case, we compared empirical designs (e.g. random and/or clustered) to local and on-average D-optimal ones, assuming they both outperform the empirical designs. Theoretically, locally optimal designs should be the most efficient ones regarding the information gain on the processes. However, they require a good knowledge on parameters before the survey which is, in practice, rarely the case. Thus, given the usual uncertainty on populations traits and considering on-average designs appears more relevant.

### 6.1 Agroecological case study

In most studies investigation insect populations in agricultural landscapes, traps are placed according to what we call a clustered design. In practice, when a field is chosen, three spatially grouped traps are put in this field as illustrated in Fig. 1b. For this case, we compared some properties of local D-optimal, on-average D-optimal, clustered and random (i.e. where spatial locations are drawn from a binomial point process) designs. The expected value and the range of estimated parameters used for respectively local and on-average designs are given in Appendix A.1.

**Figure 1:**
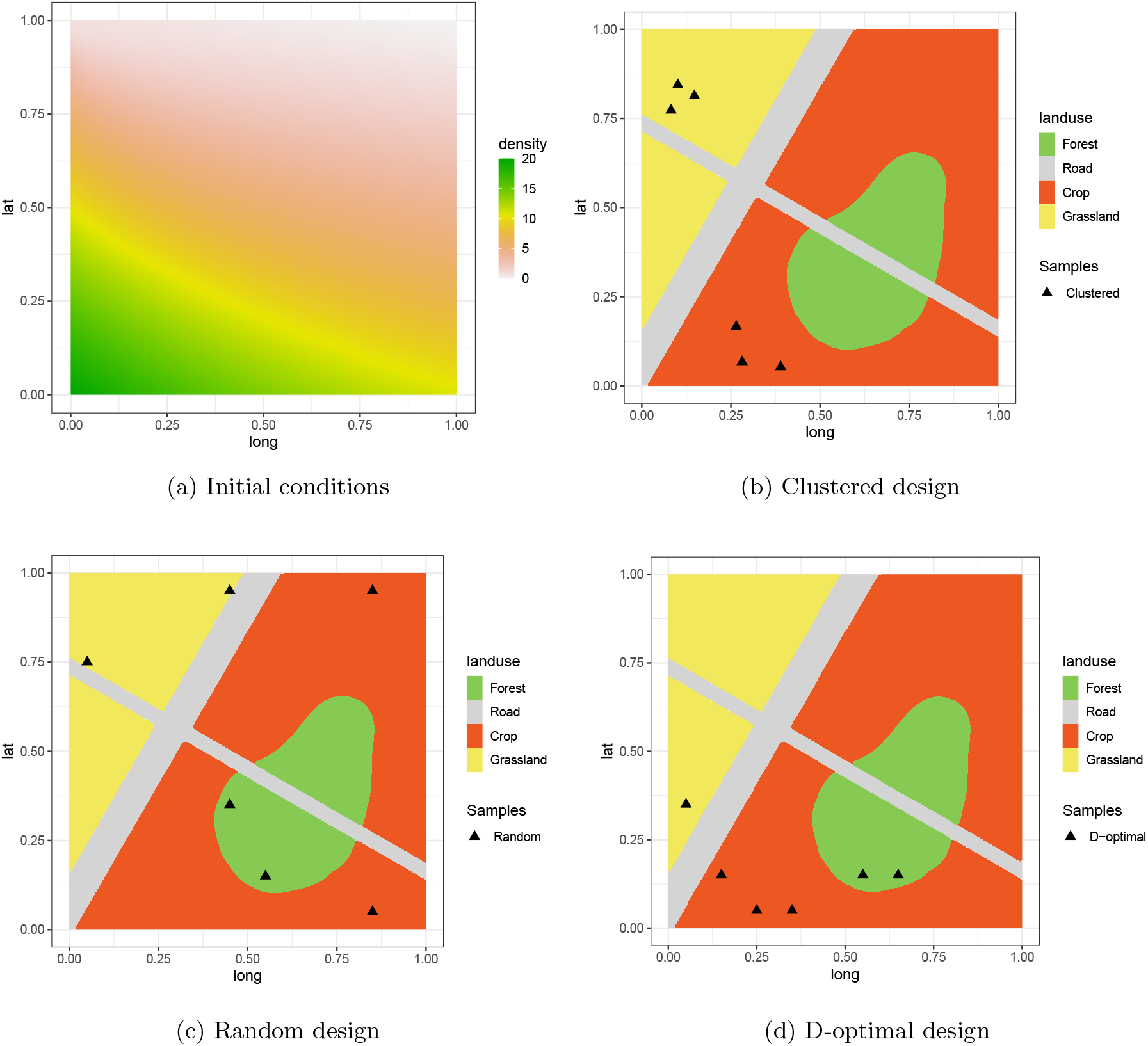
Pitfal trap placements comparisons : (a) initial population densities; (b) clustered design with 3 traps per field (pseudo-replicate) in different fields; (c) randomly positioned pitfal traps ; (d) local D-optimal design

As shown in Figure 1, the initial conditions were generated as a gradient from bottom left to top right, and the landscape contains 2 crop fields (orange), two grasslands (yellow), one region of semi-natural habitat (e.g. forest patch in green) and two crossing roads (gray).

The local D-optimal design displayed in Figure 1d concentrates the points on the bottom left corner of the landscape where the highest slope of initial conditions is. Moreover, it seems to cover all the habitats, even if unequally (except roads where the growth rate is set to 0), forming a stencil (as in numerical analysis [40]) with the added constraint to cover all the habitats.

Figure 2 shows the position of the traps in an on-average design and 95% confidence ellipses, estimated by inverting the Fisher information matrix, for the pairs of growth rates (*r*_*c*_, *r*_*s*_), (*r*_*c*_, *r*_*g*_) and (*r*_*s*_, *r*_*g*_). Interestingly, compared to the local-design, the stencil shape seems less apparent here as we see four points align within the initial condition density but two points are farther away and distributed between two different habitats. As expected, the confidence ellipses surface are smaller for all pairs of parameters for the on-average D-optimal design compared to the cluster one (Fig. 2c-d & Appendix A.2 for those related to *β, µ* and *D*). The uncertainty associated with semi-natural growth rate *r*_*s*_ is severely reduced which is logical given the clustered design did not account for that habitat. In fact, the optimal design suggests that only one point within this habitat drastically reduces the variance of its estimates.

**Figure 2:**
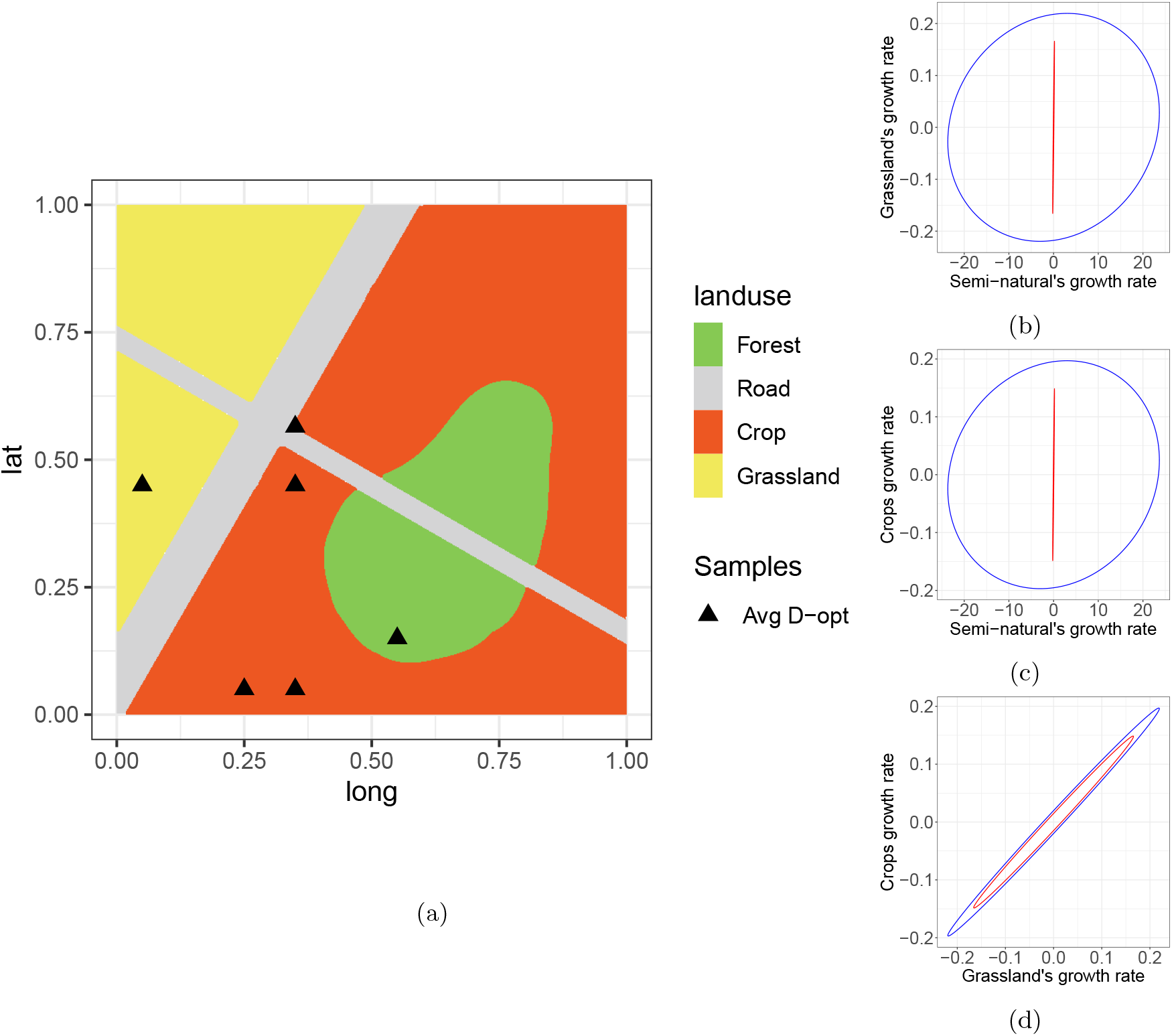
On-average exact D-optimal design : (a) position of pitfal traps in case of on-average D-optimal exact design ; (b-d) 95% confidence ellipses for pair of growth rates, estimated by inverting the Fisher Information Matrix for *θ*, for the on-average optimal design (red) and the clustered design (blue) seen in fig. 1.

The comparison of the efficiency of the designs in relation to the tour length is presented in figure 3 for both local and on-average designs. Random designs are represented by black circles (there are a hundred of them), the clustered design by a blue dot, the D-optimal design by a green dot and if at least one red dot exist, it is a random sampling that is pareto optimal for at least one dimension. One can see that the clustered design is, of course, less statistically efficient than the D-optimal but also that it has a longer tour. In fact, the D-optimal designs here are dominant both in statistical and tour length. Several simulations of random designs show that it is not always the case but the optimal designs are still among the shortest (data not shown). One can also note that most random designs are more D-efficient than the clustered design. This is probably because they either place traps within semi-natural habitat and/or simply cover more ground, with a longer tour length. As in this study we drew only one clustered empirical design, the solid comparison between random and clustered sampling strategies, which was not the scope of this work, would require more simulations.

**Figure 3:**
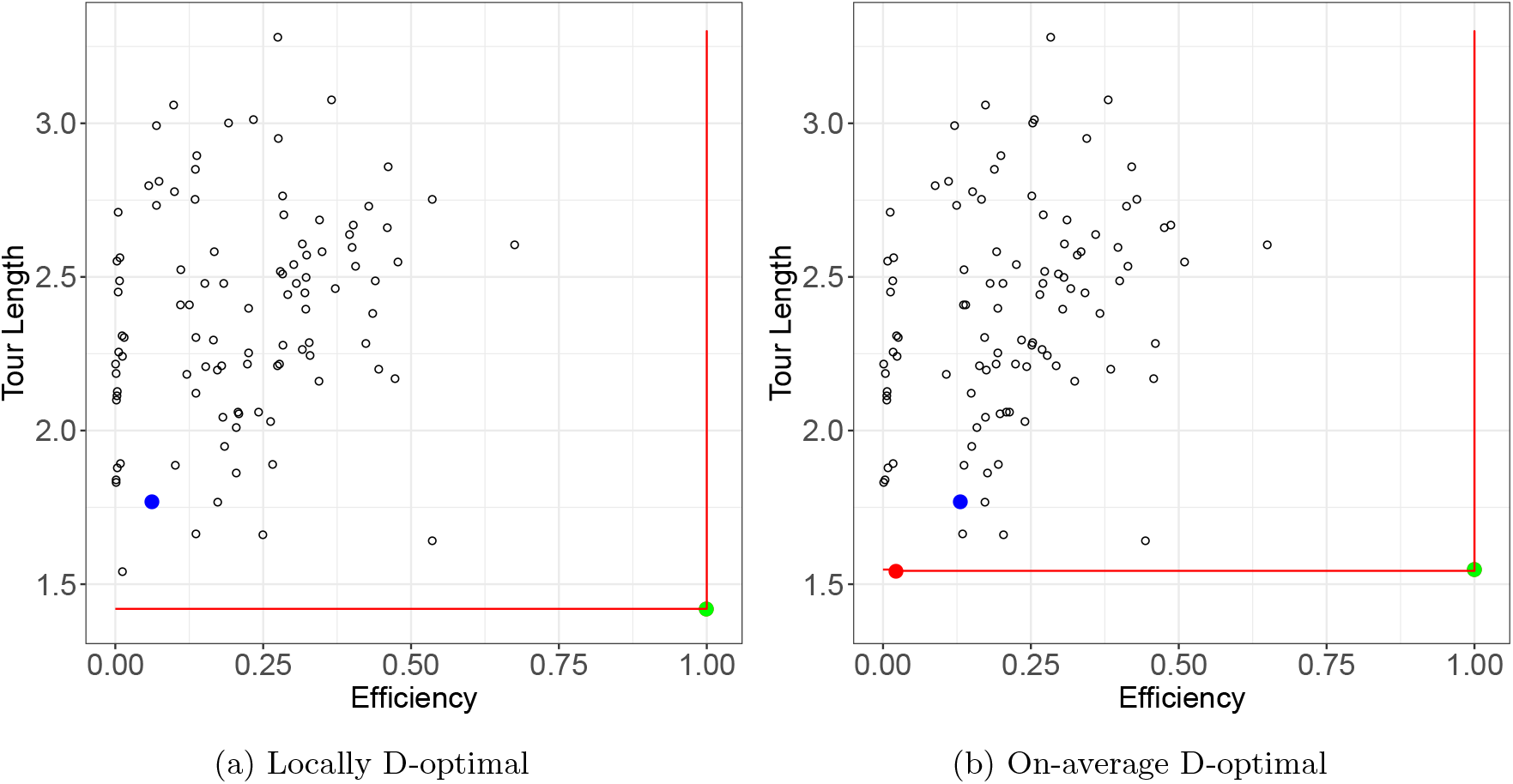
Linear approximation of the Pareto front : (a) for the local design; (b) for the on-average design. Each point represents the D-efficiency (x) and tour length (y) of a design. Black circles represent 100 random samplings (the same set is projected on each panel), blue dots represents the clustered sampling, green dots are optimal and red lines and dots form the linear approximation to the pareto fronts. All the designs D-efficiencies are evaluated locally (a) or on-average (b).

### 6.2 Conservation biology case study

In this example the design problem consists in searching *n* = 8 aerial transects, of different lengths, chosen among N=32 reported in [29]. The expected values and the range of parameters *θ* are given in Appendix A.1.

As shown in figure 4a the monitored area has less high quality habitats (green) than average and low quality ones (grey). The original transect coverage was extremely dense (Fig. 4b) and it appears natural to reduce it, for instance by taking only 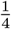 of the possible transects. As short transects are frequent in the original empirical distribution of transects, there are a few chances of randomly choosing the longer ones (Fig. 4c). When defining a local D-optimal design with a ‘low growth and dispersal’ scenario, the longer and more informative transects appear to be favored (Fig. 4d).

**Figure 4:**
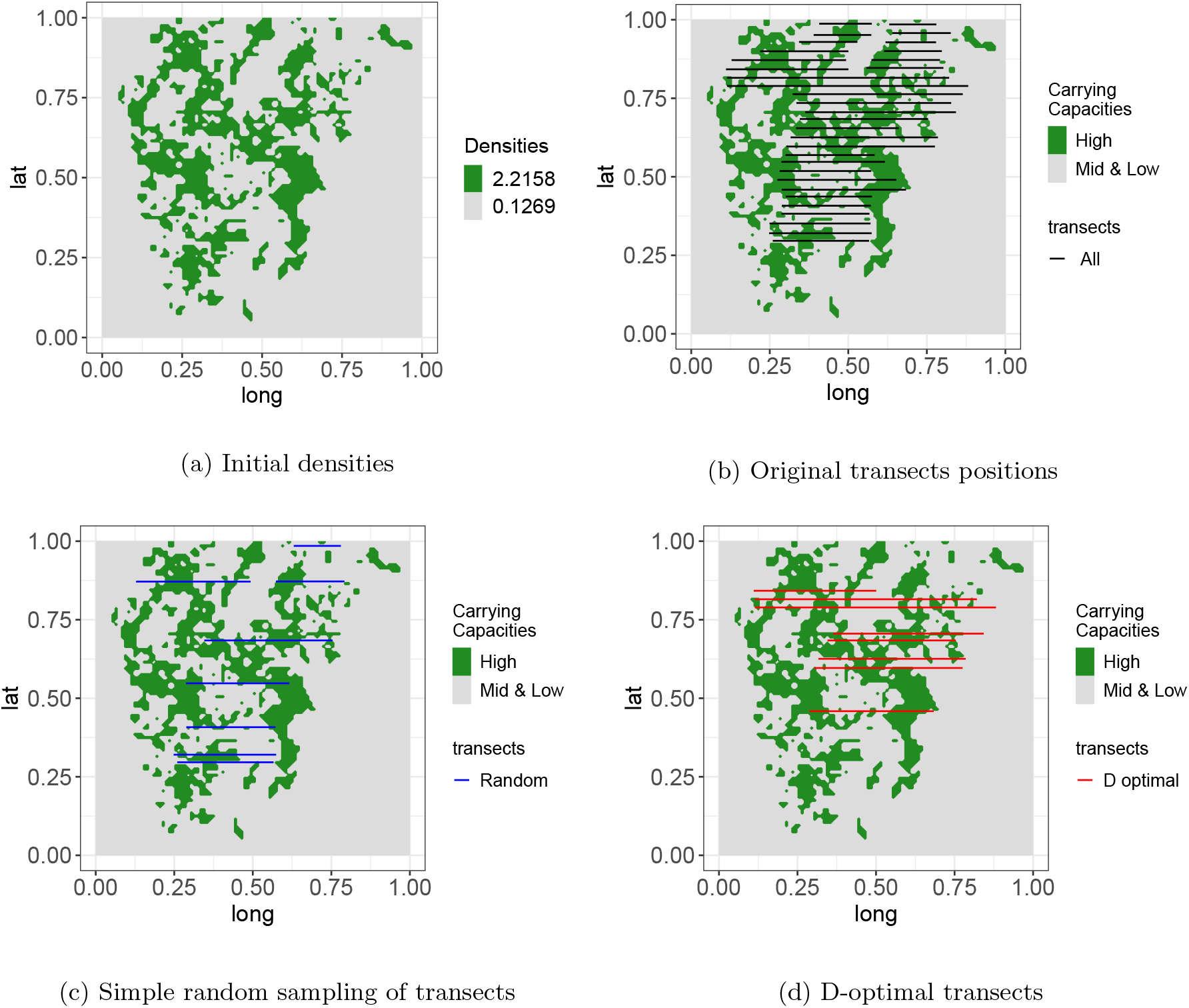
Transects comparisons: In the original study, high initial densities were linked with a high carrying capacity while mid and low densities were linked with another carrying capacity (see also Appendix A.1). We can see here (a) the initial population densities (b) then all 32 original transects (in black) over the carrying capacities ; (c) some random subset of 8 of the transects (in blue) ; (d) finally a locally D-optimal design of 8 transects for *θ*_*min*_ (i.e. ‘low growth and dispersal’ scenario) (in red)

The on-average D-optimal design (5b) appears to be similar to the local design (Fig. 4d). It seems to favor longer transects and, in that case, only one transect on the southern part of the map. The spatial extent of the optimal transects is smaller than most random transects but, as it favors longer transects, the tour length, forcibly passing by transects, is actually longer than most (Fig. 5d). As expected, D-optimality reduces the confidence ellipse of estimated parameters compared to a random design (Fig. 5c). The variance reduction appears to be greater for the diffusion coefficient *D* than for the growth rate *b*. Finally, the Pareto front displayed in Figure 5d to assess the compromise between design efficiency and tour length, shows that the on-average D-optimal transects dominate the D-efficiency but is here the worst choice in term of tour length. This means that there are multiple point on the Pareto front that could be candidate if a compromise has to be chosen between D-efficiency and tour length.

**Figure 5:**
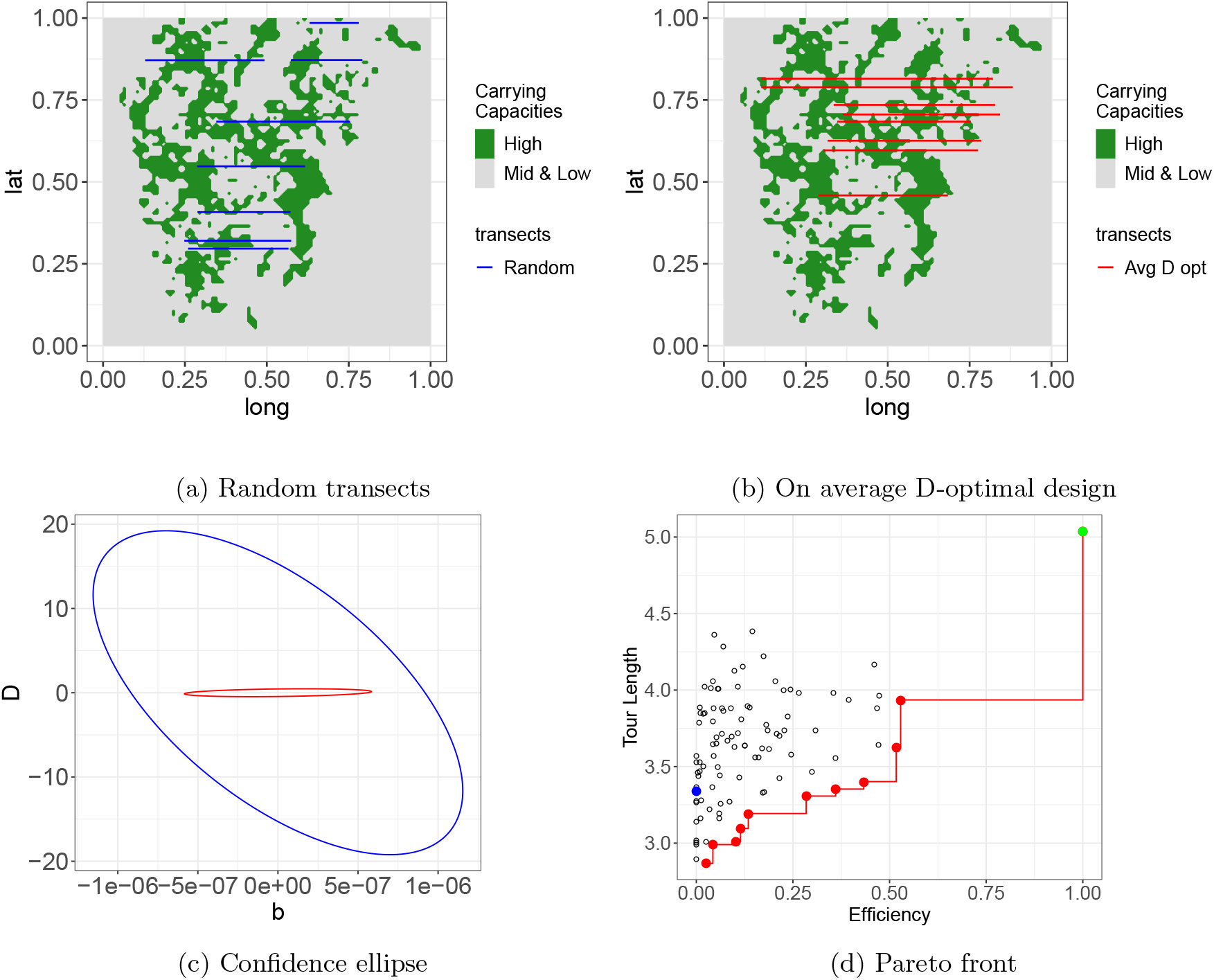
On-average exact D-optimal design : (a) random transects; (b) on-average D-optimal ; (c) 95% confidence ellipse of b and D, blue is for the random design figured in (a) and red for the optimal design, both evaluated at *θ*_*min*_ and, finally, (d) a linear approximation of the Pareto front with D-efficiency evaluated at *θ*_*min*_ (color coding and shapes are the same as in figure 3)

## 7 Discussion

In this study we addressed the question of where to make observations within a large spatial domain to efficiently capture the changes in space and time of a population described by a reaction-diffusion model. By considering two example systems, i.e. the dynamics of carabid beetles in agricultural landscapes and wild horses in the Australian Alps, we showed that the use of the optimal design of experiments framework can be useful to find optimal locations for trapping or counting individuals that maximize the information on the population for parameter estimation. In particular, we were able to produce spatial designs that outperform both those used in the initial studies and random designs. More precisely we first obtained local D-optimal designs that assume that the expected values of parameters are known before planning data collection. Then, as this situation is generally unlikely to occur in the population ecology, we investigated the situations when only bounds around population parameters are known considering on-average D-optimal designs. While local designs appeared to be obviously the most efficient for minimizing the variance of parameters estimates, our results points out the usefulness of optimal design without strong *a priori* to think spatial monitoring so it will enable efficient model fitting.

Although the question of optimized survey in ecology or epidemiology is still seldom considered, it has already been addressed in several studies using various frameworks. For instance, [8] proposed a framework that produces designs that minimizes prediction uncertainty, [9] calculated optimal observation times for botanical epidemiology experiments by maximizing the Kullback-Leibler divergence from the posterior to the prior, or [41] who proposed a strategy for choosing insect traps locations by minimizing uncertainty in neural networks outputs. Here, we considered the optimal design of experiments framework that is briefly introduced in part 3. While its use is common for solving experimental design issues in environmental sciences or systems biology [42], it is poorly considered in ecology. The monitoring of population on large spatial domain is actually similar to the problem of optimal sensors placement for spatially distributed systems [3].

Despite the recent development of devices for increasing ecological monitoring, in practice the survey of population in large spatial domains remains challenging and a major bottleneck to the study of how populations change in space and time and the design of efficient management strategies [43, 44]. As stated above, statistical frameworks developed to optimize how and where one should make observations can be used to design more efficient population surveys. This model-based design of experiments can also take into account several practical constraints. In our study we chose to consider the tour length between traps for collecting carabid bettles and aerial transects from which wild horses are enumerated. Then we used a Pareto front to asses how designs efficiency changes with the tour length. As illustrated in Fig. 5d, it is possible to find good candidate that maximizes design efficiency while minimizing tour length so one can choose those that fit the compromises that are acceptable. For the agroecology example, we found that the best design was also minimizing the tour length. This may be explained by the smaller spatial extent of the domain compared to the wild horse case. In fact, it is likely that increasing the spatial domain would create a more progressive Pareto front between design efficiency and tour length. Depending on the considered problem, other constraints could be included in the approach to find designs that are statistically efficient and minimized the constraints. For instance one could imagine several travellers and seek for the strategy that minimizes the cost of the survey, considering both human and transport costs. While, in our opinion, finding monitoring designs that provide maximum information on population spread is an important question on which the community of modellers in ecology should concentrate more, integrating realistic constraints in model-based design is likely to catch the interest of field ecologists and practitioners. Moreover, in our two study cases we focused on the spatial aspect of monitoring assuming that the only degree of freedom is the survey designs was the locations of observations whereas the time of observations was fixed. This assumption could be relaxed and both temporal and spatio-temporal D-optimal designs may be obtained in further studies. Optimal times at which observations should be done may be obtained by solving local or on-average D-optimality problems before the survey starts. In addition, if applicable, the problem can also be tackled sequentially as an alternative to the on-average design. In this case, optimal groups of observations are proposed, over time, after having performed a parameter estimation on already acquired data, this approach is know as a sequential design [45].

Apart the theoretical background, the main obstacle to a widespread usage of optimal design of experiments in spatially explicit systems is the computational cost. Using usual algorithms and computing strategies (i.e. memoization and parallelization), in this study we were able to solve local and an-average D-optimality problems on reaction-diffusion models with a reasonable amount of time. However, increasing the complexity of the problem, for instance by adding more practical or economic constraints or the temporal dimension of sampling, may require more advanced computing methods and grid computing. Structural and practical identifiability issues may also arise and make the problem even more complex. Optimal design is useful for maximizing the information gain from data collection, however the modeller has to ensure that enough information is available so the parameters can be determined. In this study, before seeking for an optimal design, we assessed the practical identifiability of the model by multiple start of a numerical local approach [24] within each relevant domain Θ.

The issue of initial conditions is well known by mathematicians and modellers working with dynamic models. In some conditions, it is possible to estimate initial conditions from monitoring data. Nevertheless, in most studies one has to make strong hypothesis and find a way to fix it before performing a statistical inference, or seeking an optimal design. Regarding spatial ecology, the most frequent approach is perhaps to use past (or early data) with a regression model to someway estimate the distribution of population density at the beginning of the study. In our study, we did not include the initial conditions in the designs problems. It is very likely that optimal designs are dependent on the initial conditions. This may be assessed by augmenting the problem and exploring how changes in the initial conditions impact optimal-designs. Yet, augmenting the problem for taking care of initial conditions has its limits. To cite in extenso [46], it is “[a] very prospective direction [where] the infinite dimensional nature of the resulting parameter space is inherently associated with the ill-posedness which means that even low noise in the data may make the estimates extremely unstable.”. The reader could refer to [47, 48] for further information.

In this article we focus on the D criterion but there exist different criteria (e.g. different functions *ϕ*(.)) that can be applied to the information matrix. For example, one can seek to minimize the average variance of the estimates by minimizing the negative of the trace of the inverse of the Fisher information matrix (i.e. A-optimality). Here, we focused on the D criterion because (i) through the equivalence theorem [26], we know that main classical criteria are linked in some ways (e.g. *A* and *D*), (ii) the D criterion is invariant by reparameterization (e.g. helpful to control parameters positivity) and (iii) there exist variants of the D criterion that are useful for a variety of purposes. For instance, if one is interested in model selection, the *D*_*s*_ criterion enable to focus the problem on the parameters of interest versus ‘nuisance’ parameters, hence being a a canonical criterion in nested model selection [49]. For more general purpose model selection in case of gaussian observations, the generalized DT criterion [50] is a criterion that can be described as a normalized arithmetic mean of *D* criterions and *L*_2_ distance between predictions, handling both optimal parameters estimation and model selection.

To finish with, the purpose of this article was to push the discussion on the statistical efficiency of experimental designs linked with spatial ecological models, as question that is yet seldom expressed this way. The main goal of a field ecologist is to sample ‘as much as possible’ within the constraints of its budgets (monetary and manpower), but this strategy can either fail to capture the essential information on the population or be very expensive. In addition to the numerous studies that had demonstrated that formalizing ecological processes into mathematical models and using them to analyze empirical data offers a mean for improving our understanding of populations spread, we hope this work points out that the difference between optimal and non-optimal monitoring can be significant in term of information gained, and thus that model-based design of ecological survey is a promising path that could also contribute to reduce environmental costs of samples collection, storage and processing.

## Authors’contributions

All authors designed the numerical experiments and the overall framework. ML provided insight on ecological modeling, KAC provided fruitfull guidance on statistics, NP implemented the framework, ran the numerical experiments and provided focus on optimal design. All authors drafted, managed, reviewed and approved the manuscript.

## Acknowledgements

The authors thank Jean-Sébastien Pierre, Jacques Baudry and Solenn Stoeckel for useful discussions about this work. The authors want to thank especially Professor Darius Uciński for a brief but very supportive discussion about an early version of this work.

## A Appendix

### A.1 Use case’s parameters

A.1 Use case's parameters
| Parameters | Original values | Explored min | Explored max | units | source |
| --- | --- | --- | --- | --- | --- |
| $D$ | 76.1 | 19.025 | 304.4 | $\text{m}^2 \cdot \text{day}^{-1}$ | [1] |
| $\beta$ | 0.123 | 0.111 | 0.135 | — | [1] |
| $\mu$ | 0.210 | 0.115 | 0.42 | $\text{day}^{-1}$ | [1] |
| $r_s$ | 0.155 | 0.144 | 0.166 | $\text{day}^{-1}$ | [1] |
| $r_g$ | 0.304 | 0.293 | 0.315 | $\text{day}^{-1}$ | [1] |
| $r_c$ | 0.385 | 0.374 | 0.396 | $\text{day}^{-1}$ | [1] |
| $r_r$ | 0 | — | — | $\text{day}^{-1}$ | [1] |
| $\eta$ | $10^{-4}$ | — | — | — | [1] |
| $\tau$ | 5 | — | — | day | [1] |
| $K_H$ | 2.462 | — | — | horse | [2] |
| $K_O$ | 0.141 | — | — | horse | [2] |
| $D$ | — | 2 | 30 | nondimensional | [2] |
| $b$ | — | 0.16 | 0.27 | $\text{year}^{-1}$ | [2] |
| $\sigma$ | $10^{-3}$ | — | — | $\text{horse}^2$ | arbitrarily low |

Parameters used for the use cases when searching for local and on-average optimal designs. The symbol ′− ′means ‘not applicable’ e.g. when used for explored min or max, it means those parameters were not part of the optimal design problem so they were kept at their original values. Reference [1] is [13] and reference [2] is [18].

### A.2 Supplementary confidence ellipses

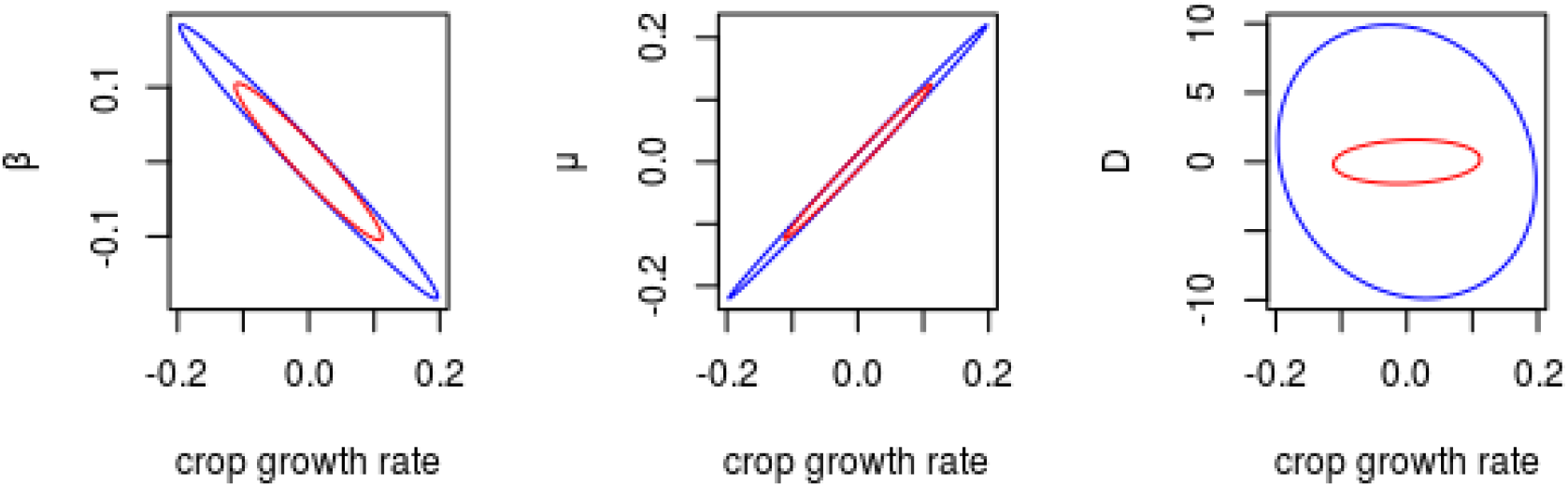

Confidence ellipses at 95% for the agroecological case study, where parameters *β, µ* and *D* are paired with the crop growth rate.

